# Disruption of the Brain-Spleen Axis Impairs Monocyte-Microglia Communication and Accelerates Disease Progression in a Model of Amyloidosis

**DOI:** 10.64898/2026.03.18.712196

**Authors:** Tommaso Croese, Miguel A. Abellanas, Hodaya Polonsky, Michal Arad, Javier M. Peralta Ramos, Yuliya Androsova, Serena Riccitelli, Sedi Medina, Francesca Palmas, Romano Strobel, Giulia Castellani, Denise Kviatcovsky, Sarah Phoebeluc-Colaiuta, Miriam Adam, Sama Murad, Hannah Partney, Daniel Kitsberg, Alexander Dieter, Tomer-Meir Salame, Alexander Brandis, Tevie Mehlman, Oded Singer, Michal Rivlin-Etzion, Simon Wiegert, Yosef Shaul, Oren Kobiler, Ofer Yizhar, Naomi Habib, Michal Schwartz

## Abstract

Alzheimer’s disease (AD) is characterized by a prolonged asymptomatic phase before cognitive decline emerges, yet the mechanisms driving symptom onset remain unclear. Here, we hypothesized that the transition from asymptomatic to symptomatic disease is linked to dysfunction of brain–immune communication. Retrograde neuronal tracing in the 5xFAD mouse model of amyloidosis revealed reduced brain–spleen connectivity at advanced disease stages. To probe the functional role of the brain–spleen axis in coping with disease, we denervated the splenic nerve at an early presymptomatic stage. This intervention accelerated cognitive decline, impaired splenic hematopoiesis, diminished monocyte recruitment to the brain, disrupted monocyte–microglia signaling networks, and reduced the transition of microglia from a homeostatic to the disease-associated (DAM) state. Conversely, enhancing splenic noradrenergic input increased hematopoiesis, restored monocyte homing to the brain, and delayed cognitive impairment. The protective role of splenic monocytes was independently validated in a retinal cytotoxic injury model, in which splenic denervation impaired post-insult retinal ganglion cell survival. Together, these findings identify an active brain–spleen circuit in regulating monocyte recruitment and establish peripheral monocytes as key drivers of microglial state transitions and disease progression.

## Introduction

Alzheimer’s disease (AD) is characterized by a prolonged clinically silent phase during which amyloid pathology, among other hallmark pathologies, accumulates in the brain long before the emergence of clinical symptoms of cognitive decline (1, 2). Although genetic and pathological studies have proposed molecular drivers of disease progression within the brain, the mechanisms that govern the transition from an asymptomatic state to overt clinical dysfunction remain poorly understood. Increasing evidence implicates the immune system as a central modulator of AD onset and progression. Genome-wide association studies have identified immune-related genes, specifically linked to microglia and monocytes, as major contributors to AD susceptibility, and functional studies have demonstrated that immune responses can both restrict and exacerbate disease pathology (3–5). Yet, the extent to which systemic immune regulation affects the timing of symptom onset remains unclear.

Within the brain, microglia have been shown to undergo dynamic state transitions during AD, shifting at early disease stages to acquire a disease-associated phenotype (DAM) that has been proposed to limit pathology and support neuronal integrity at the early disease stage (6). However, as pathology advances, microglia become exhausted or senescent, potentially compromising their protective capacity (7).

In parallel, accumulating evidence indicates that peripheral immune cells, particularly monocyte-derived macrophages, contribute to CNS protection and repair under pathological conditions (5, 8–11). In AD mouse models of amyloidosis, monocyte recruitment to the brain has been shown to enhance amyloid clearance and modulate neuroinflammation (12–15). Thus, recruitment of peripheral immune cells under disease conditions such as AD might be a key mechanism for maintaining brain health, yet their spontaneous mobilization to the brain is typically limited and tightly regulated. These findings raise a fundamental question: What governs the engagement of peripheral immunity during chronic neurodegeneration?

The brain actively communicates with peripheral immune organs through autonomic neural circuits. Sympathetic and parasympathetic pathways regulate immune cell activation, cytokine production, and hematopoiesis (16, 17). This is well exemplified by the inflammatory reflex, in which cholinergic vagal efferents of the parasympathetic nervous system sense peripheral immune activation and dampen inflammation in visceral organs by suppressing the release of pro-inflammatory cytokines. However, whether disruption of brain-to-immune signaling occurs in the course of chronic neurodegeneration, and if so, whether it contributes to disease progression, has not been directly tested.

Here, we hypothesized that communication between the brain and immune organs regulates peripheral myeloid responses required to sustain protective microglial functions during AD. Specifically, we focused on the spleen — a key site of emergency hematopoiesis and monocyte maturation — and postulated that disruption of the brain–spleen axis may limit monocyte recruitment to the diseased brain and accelerate the onset of cognitive decline by altering signaling to resident brain cells.

We found that brain–spleen connectivity declines at advanced disease stages. Partial splenic denervation, performed long before symptom onset, accelerated cognitive decline, impaired monocyte homing to the brain, and reduced microglial transition to the DAM state. Conversely, enhancing noradrenergic signaling in the spleen increased monocyte recruitment to the brain and mitigated the cognitive impairment. Further generalizing these findings we showed that spleen denervation also had a negative effect on recovery in a model of retinal insult. Together, these findings identify the brain-spleen axis as a critical regulator of neuroprotective processes within the CNS by controlling monocyte homing to the brain and shaping microglia state transitions that impact disease progression and repair.

## RESULTS

### The role of brain–spleen communication in disease manifestation in 5xFAD mice

The spleen, a crucial site for immune cell regulation and maturation (18–20), receives its innervation from the splenic nerve, which contains mainly sympathetic fibers (21, 22). These sympathetic fibers release norepinephrine, which initiates an anti-inflammatory pathway involving T cells and macrophages (23–25). Using the 5xFAD mouse model of amyloidosis, we first assessed whether spleen innervation is impaired at an advanced disease stage, when cognitive deterioration is fully manifested. To this end, we performed retrograde tracing of neurons in the brain by injecting a pseudorabies virus encoding a red-fluorescent protein (PRV-RFP) into the spleen of 5xFAD and wild-type (WT) littermates at the age of 12 months (**Fig. 1a**). We examined RFP^+^ neurons 4 days after virus injection in brain regions implicated in the regulation of this autonomic pathway, including the C1 neurons of the rostral ventrolateral medulla (RVLM), the N10 portion of the dorsal motor nucleus of the vagus, and the locus coeruleus (LC). An overall reduction in labeled neurons was observed across all regions in 5xFAD mice compared with WT controls, indicating diminished neuronal connections between the spleen and CNS (**Fig. 1b-f**).

**Fig. 1:**
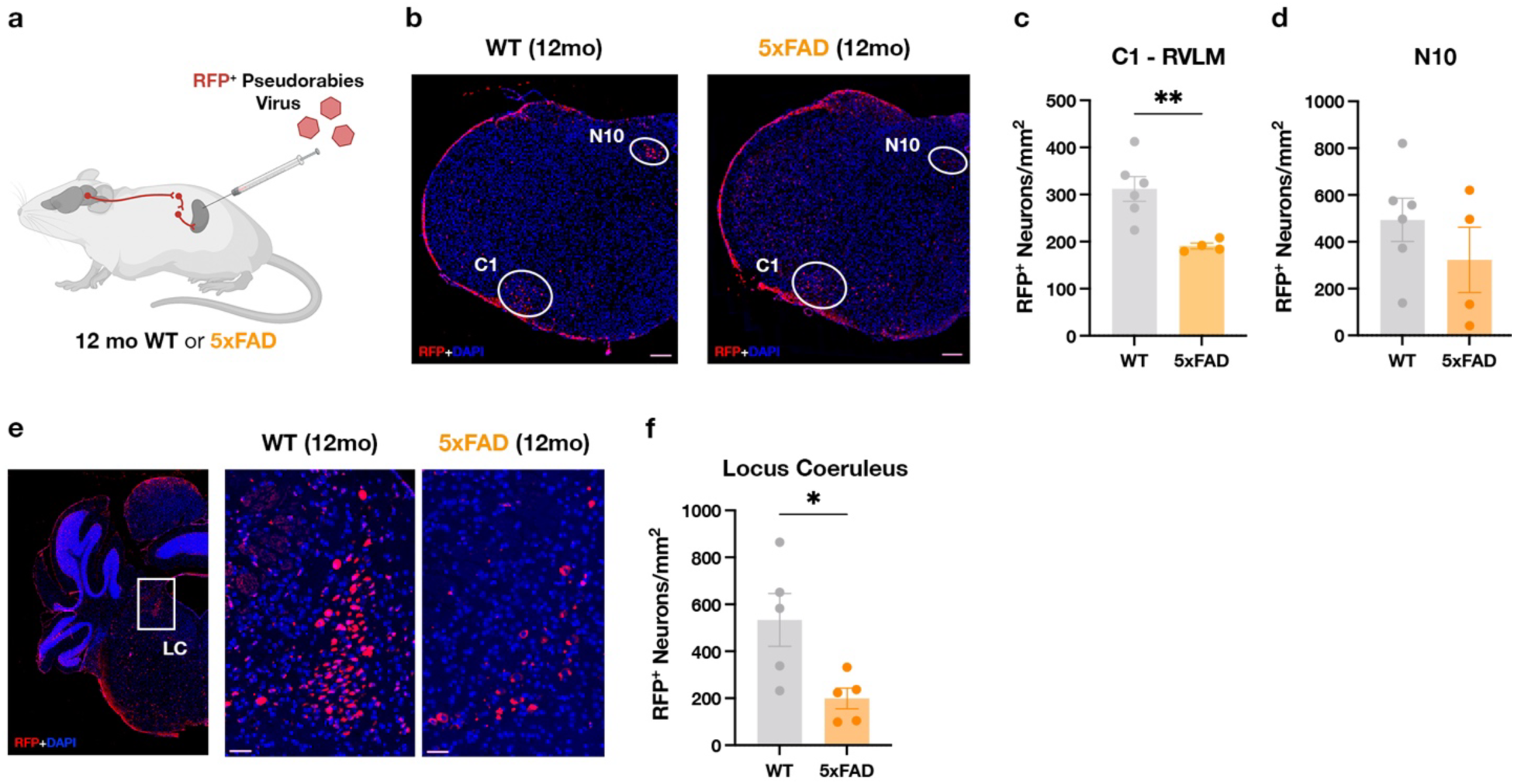
Brain-Spleen connectivity is partially lost in the 5xFAD model. **a.** Pseudorabies virus (PRV) encoding red fluorescent protein (RFP) was injected into the spleens of 12-month-old 5xFAD (n= 2M; n= 2F) or WT littermates (n= 2M; n= 4F). After 4 days, the presence of RFP was analyzed by immunofluorescence in several brain regions. **b**, Representative images showing RFP expression in the N10 portion of the dorsal motor nucleus of the vagus nerve and the C1 neurons of the rostral ventrolateral medulla (RVLM). Scale bar: 200 μm. **c**, Density of RFP^+^ neurons among the C1 neurons of the RVLM. **d**, Density of RFP^+^ neurons in the N10 portion of the dorsal motor nucleus of the vagus nerve. **e**, Representative images showing RFP expression in the locus coeruleus (LC). Scale bar: 50 μm. **f,** Density of RFP^+^ neurons in the LC. Unpaired t-test. In all panels, data are represented as mean ±SEM. *p < 0.05, **p < 0.01.

### Splenic denervation impairs extramedullary hematopoiesis and alters monocyte profiles and migratory capacity

To determine if reduced brain–spleen communication affects disease progression, we experimentally interfered with the brain-spleen communication long before symptom onset to test whether this intervention would accelerate disease manifestation. We first established a protocol of partial denervation of the brain-spleen communication in WT mice. To this end, we ethanolized the splenic nerve plexus (26, 27). We then injected PRV-RFP into the spleen 2 weeks post-denervation (**Fig. 2a**) and quantified the number of retrogradely labeled neurons in the LC. Injection of PRV-RFP into the spleen of denervated WT mice revealed a marked reduction in spleen-projecting neurons within the LC compared to sham-operated controls **(Fig. 2b)**, while the total number of TH^+^ neurons in the LC remained unchanged **(Extended Data Fig. 1a).** We further analyzed neurotransmitters such as serotonin (5-HT), acetylcholine (ACh), norepinephrine (NE), and dopamine (DA) in the spleens of WT mice 4 weeks post-denervation. NE levels were significantly lower in the denervated spleens compared to sham controls (**Fig. 2c**), while plasma NE remained unchanged (**Extended Data Fig. 1b**). Other neurotransmitters were unchanged in the spleens after denervation (**Extended Data Fig. 1c**). Additionally, there was a decrease in catecholaminergic (TH^+^) fibers in the denervated spleens (**Extended Data Fig. 1d**). These results confirm the loss of noradrenergic input to the spleen following the denervation procedure. Having established the splenic denervation model, we next assessed its impact on immune cell composition and function. Mass cytometry (CyTOF) analysis of WT and 5xFAD mice 4 weeks post-denervation revealed no significant changes in the frequency of major immune cell subsets (**Extended Data Fig. 1e, f**).

**Fig. 2:**
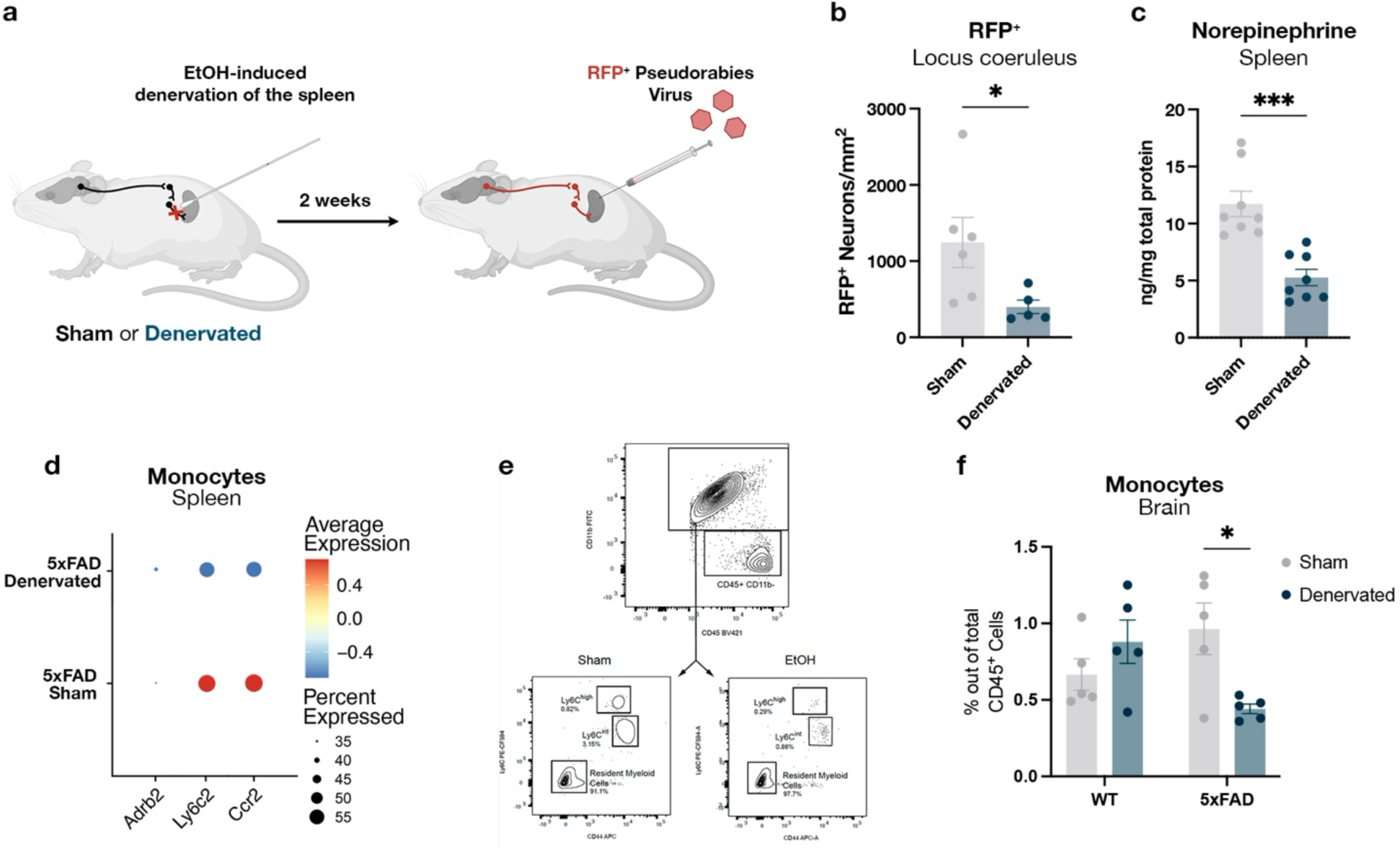
Surgical denervation of the splenic nerve impairs monocyte migratory capacity to the brain. **a**, Scheme depicting neural tracing assay by RFP-encoding pseudorabies virus injection 2 weeks after denervation of the spleen. **b**, Quantification of RFP^+^ neurons in the locus coeruleus of sham (n= 3M; n= 3F) and denervated (n= 2M; n= 3F) animals. **c**, Levels of norepinephrine in the spleen comparing sham (n= 4M; n= 4F) vs. denervated (n= 4M; n= 4F) animals 4 weeks after denervation. **d**, Dot-plot showing scaled expression of selected genes in scRNAseq analysis of monocytes from the spleen of denervated (n= 2M; n= 2F) or sham-operated (n= 2M; n= 2F) 5xFAD mice. **e**, Gating strategy of flow cytometry analysis of monocytes from the brain of sham or denervated WT and 5xFAD mice. **f**, Monocytes detected in the brains of sham or denervated WT (n= 6M; n= 4F) and 5xFAD (n=5M; n= 5F) animals. Unpaired t-test or two-way ANOVA followed by Sidak’s multiple comparison test. In all panels, data are represented as mean ±SEM. *p < 0.05, ***p < 0.001.

The spleen is a preferred site for extramedullary hematopoiesis (28), supporting bone marrow hematopoiesis by boosting myeloid cell differentiation in inflammatory contexts (29). Previous studies showed that sympathetic innervation regulates extramedullary myelopoiesis in the spleen (30) and facilitates the release of Ly6C^high^ monocytes (31), a subset of myeloid cells with high migratory potential to tissues (32–34). Therefore, we focused on how partial splenic denervation impacts myelopoiesis and monocyte profiles. Using a targeted flow cytometry panel, we analyzed the hematopoietic stem cells (HSC) present in the spleen of 5xFAD mice following denervation (**Extended Data Fig. 1g**). While the overall numbers of hematopoietic stem and progenitor cells (Lineage^-^Sca1^+^cKit^+^, LSK) remained unaltered (**Extended Data Fig. 1h**), we observed a reduction in short-term (ST)-HSC (LSK CD150^-^CD48^-^) in the spleens of 4-month-old denervated mice compared to sham-operated animals, 2 weeks after denervation (**Extended Data Fig. 1i**). Our results showed that splenic denervation led to a loss of a subset of ST-HSC, which constitute a fundamental stage in the differentiation of myeloid cells (35).

Splenic monocytes were further profiled by scRNA-seq 4 weeks after denervation. Among all immune cell subsets, *Adrb2*, which encodes the β2-adrenergic receptor (the main norepinephrine receptor on immune cells), was predominantly expressed by monocytes (**Extended Data Fig. 1j**), and its expression by monocytes was reduced in the denervated mice (**Fig. 2d**). We also detected reduced expression of *Ccr2* and *Ly6c2*, both required for monocyte homing to the brain (**Fig. 2d**). Consistent with these transcriptional changes, flow cytometry and CyTOF analyses revealed significantly lower numbers of monocytes in the brains of 4–5-month-old 5xFAD mice at both 2 and 4 weeks post-denervation compared to sham-operated animals (**Fig. 2e, f**; **Extended Data Fig. 2k-n**). Together, these findings indicated that disrupted brain–spleen communication altered splenic extramedullary hematopoiesis, modified monocyte transcriptional profiles, and reduced monocyte recruitment to the CNS in diseased animals.

### Splenic denervation accelerates symptom onset in 5xFAD mice

To determine whether splenic denervation, which resulted in reduced monocyte levels in the brain, is sufficient to affect disease progression, we assessed cognitive performance 5 to 8 weeks after surgery, allowing sufficient time for recovery. We assessed working, spatial, and recognition memory performance, previously shown to decline in the 5xFAD model (36), using the Radial Arm Water Maze (RAWM) and the Novel Object Recognition (NOR) tests (**Fig. 3a**). Spleen denervation at this pre-symptomatic stage resulted in cognitive decline in the 5xFAD mice, while age-matched sham-operated 5xFAD mice remained cognitively intact in both tests (**Fig. 3b, c**). Cognitive performance in WT mice was unaffected by denervation, consistent with prior reports that healthy brain function is not monocyte-dependent (15). Motor function, assessed in the open-field test, was also unaffected (**Extended Data Fig 2a**).

**Fig. 3:**
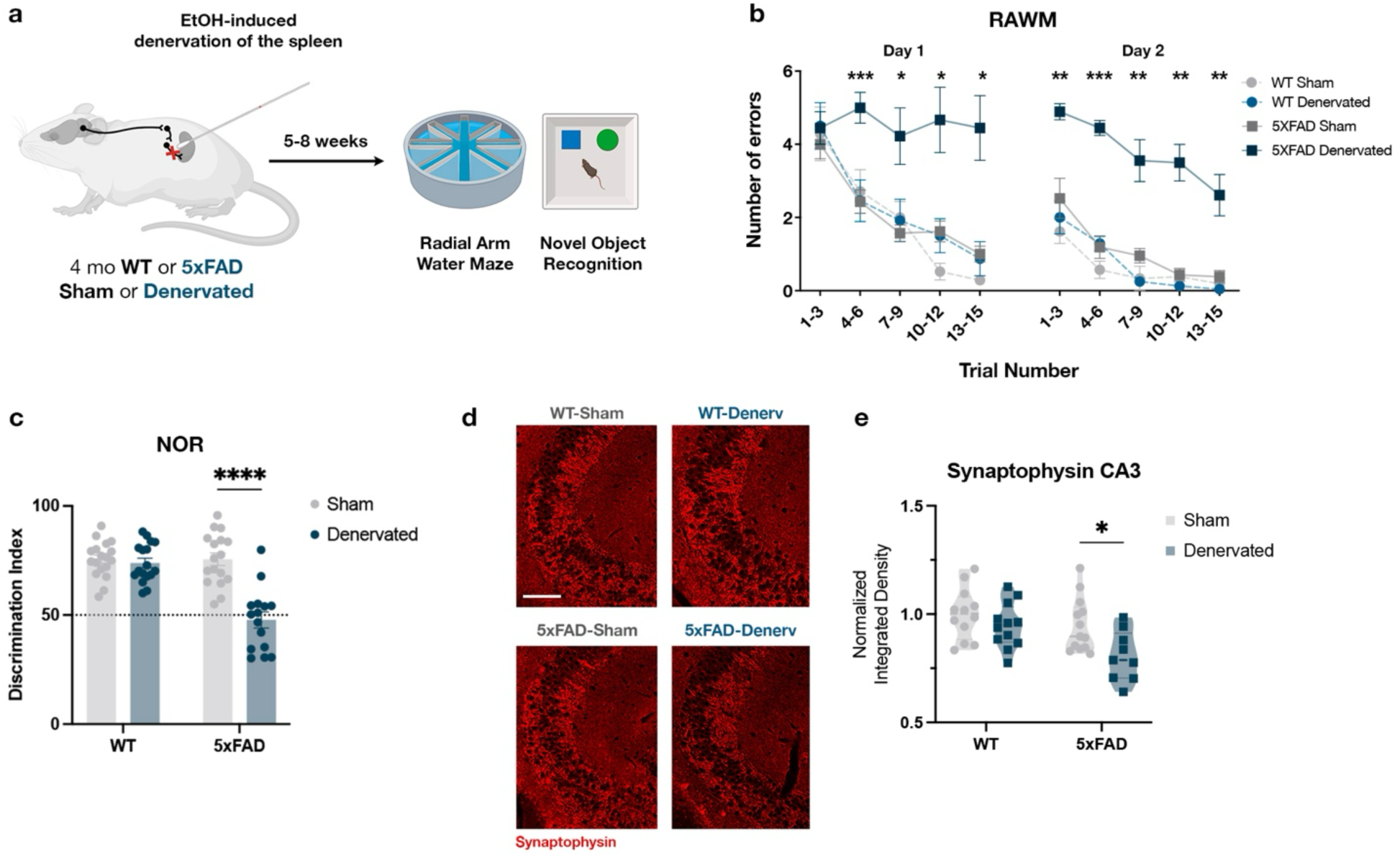
Surgical splenic nerve denervation accelerates disease onset in 5xFAD mice. **a**, Schematic representation of the cognitive tests performed 5-8 weeks after surgical denervation of the spleen of pre-symptomatic 4-month-old 5xFAD (n= 15M; n= 20F) or WT (n= 17M; n= 15F) mice. **b**, Number of errors in the radial arm water maze (RAWM) test; each point represents the average of three consecutive trials. **c**, Discrimination index in the novel object recognition (NOR) test, calculated as the percentage of time spent with the novel object over the total exploration time of the two objects. **d**, Representative immunofluorescence of synaptophysin in the CA3 region of the hippocampus. Scale bar: 100 μm. **e**, Quantification of the expression of synaptophysin by immunofluorescence in the CA3 in the hippocampus. Two-way ANOVA followed by Sidak’s multiple comparison test. In all panels, data are represented as mean ±SEM. *p < 0.05, **p < 0.01, ***p < 0.001.

Next, we examined pathological changes in the brain following splenic denervation. Synaptophysin, a marker of synaptic integrity, was significantly reduced in the CA3 of denervated 5xFAD mice compared with sham controls, with no effect observed in WT mice (**Fig. 3d, e**). As expected at this early disease stage, amyloid plaques were already present in small numbers in the hippocampus of 5xFAD mice (**Extended Data Fig. 2b, c**), while astrogliosis was not yet apparent compared with WT mice, and denervation did not alter GFAP immunoreactivity (**Extended Data Fig. 2d-f**).

Increased microglial reactivity is an early event in AD progression (37). Consequently, microgliosis, measured by Iba1 immunohistochemistry, was already observed in the hippocampal CA3 region and the dentate gyrus (DG) of 5xFAD mice, and was unaffected by denervation (**Extended Data Fig. 3g-i**). Thus, splenic denervation at a presymptomatic stage exacerbates synaptic dysfunction and accelerates cognitive decline in 5xFAD mice at an early stage when overt amyloid and glial pathology is still modest.

### Reduced monocyte-homing to the brain following splenic denervation limits microglia transition to the DAM state in 5xFAD mice

The cognitive deterioration observed after spleen denervation prompted us to investigate mechanisms through which reduced monocyte numbers in the brain might accelerate disease progression. We focused on microglia, given their central role in AD (38–40). At early disease stages, microglia acquire the DAM activation state (6), found in close vicinity to amyloid-beta plaques (41). Using CyTOF, we found significantly lower abundance of microglia with the DAM signature in 5xFAD mice 4 weeks post-denervation compared with sham-operated controls (**Fig. 4a-c**).

**Fig. 4:**
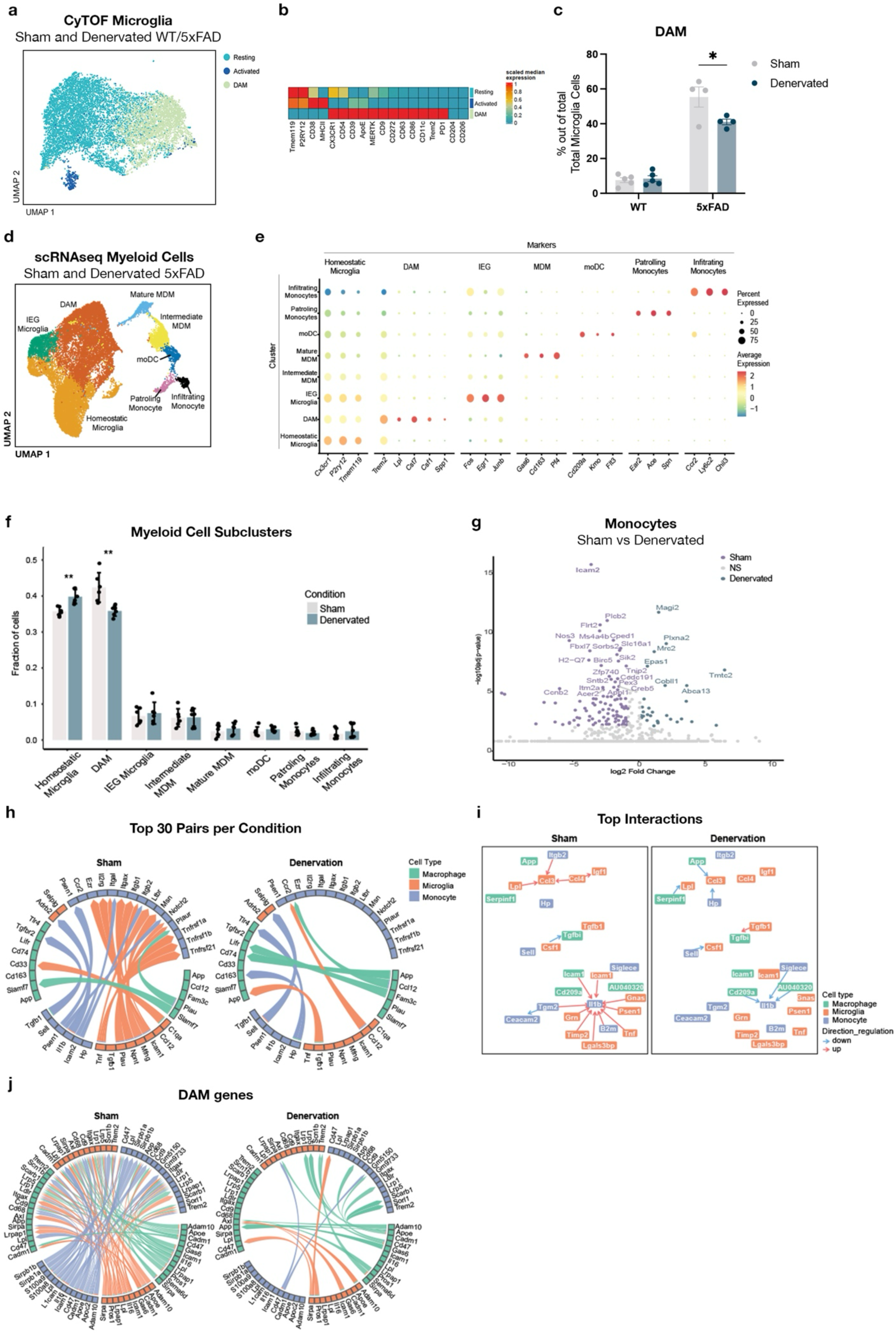
Denervation of the spleen alters monocyte-microglia communication and acquisition of DAM state by microglia. **a**, Dimensional reduction by UMAP of mass cytometry (CyTOF) data of microglia in the brain of sham or denervated WT (n= 10M) or 5xFAD (n= 8M) mice. **b**, Heatmap showing markers studied by CyTOF to annotate microglial subsets. **c**, Abundance of DAM as determined by CyTOF. **d**, UMAP embedding of 19,894 single-cell RNA profiles, color-coded by microglia subtypes, monocyte subtypes, and MDM (Monocyte-Derived Macrophages) subtypes. **e**, Mean expression level of markers used for annotation of the clusters. **f**, Proportions of the microglial, monocytes, and MDM subtypes in sham and denervated 5xFAD mice. **g**, Differentially expressed genes in monocytes (log2 FC > 0.5, adjusted p-value (FDR) < 0.05). **h**, Top 30 ligand-receptor pairs per condition. Bottom blocks of the circle: ligands expressed by sender cell types. Top blocks of the circle: receptors expressed by receiver cell types. Arrows between blocks: direction of each interaction. **i**, A network of the top 150 ligand–target interactions per condition. Arrows between blocks: direction of each interaction, from ligand in the sender cell to the target gene downstream of the receptor in the receiver cell. Color of arrow: direction of regulation up/down. Color of blocks: cell type. **j**, DAM signature genes (from Keren-Shaul et al., 2017) across conditions. Bottom blocks of the circle: ligands expressed by sender cell types. Top blocks of the circle: receptors expressed by receiver cell types. Arrows between blocks: direction of each interaction.

To further characterize the microglial profiles, we performed scRNA-seq of FACS-sorted whole-brain-enriched myeloid cells (CD45^+^ CD11b^+^) enriched from whole brains of denervated 5xFAD mice compared to sham-treated 5xFAD mice. Using an unsupervised clustering approach, we identified distinct groups of immune and microglial cell types in the brain, including monocytes and macrophages (**Extended Data Fig. 3a-g**). Sub-clustering analysis of the microglial cells classified them into three distinct states: homeostatic (expressing *Cx3cr1* and *P2ry12*), an activated state (expressing immediate early genes (IEGs), e.g., *Fos* and *Junb*), and the DAM state (expressing *Lpl*, *Cst7*, and *Trem2*) (6, 42) (**Fig. 4d, e**). In line with CyTOF data, denervation in 5xFAD mice led to a significant reduction in microglia in the DAM state compared to sham-operated 5xFAD controls, and was accompanied by an increase in homeostatic microglia (**Fig. 4f**), suggesting that denervation impaired microglial transition to the DAM state.

Differential expression analysis comparing monocytes between sham and denervated 5xFAD mice, revealed downregulation in denervated mice of *Icam2*, which encodes a critical transmembrane glycoprotein that facilitates leukocyte adhesive interactions and extravasation across the vascular endothelium (**Fig. 4g**). These findings are consistent with the downregulation of trafficking-associated genes we observed in splenic monocytes (**Fig. 2d)** and the reduced number of monocytes homing to the brain following denervation (**Fig. 2f**), indicating that denervation impaired the migratory capacity of monocytes and homing into the AD brain.

To test whether the reduced level of microglia with the DAM signature was linked to monocyte homing to the brain, we blocked the CCR2-CCL2 axis in non-denervated 5xFAD mice. We found that the CCR2 blockade effectively reduced monocyte and macrophage homing to the brain, resulting in an increase in the abundance of resident microglial cells (**Extended Data Fig. 4a-c**), and a decrease in the abundance of microglia with an activated/DAM signature, as measured by CyTOF (**Extended Data Fig. 4d-f**).

### Impaired monocyte-microglia signaling following splenic-denervation in 5xFAD mice

The link between impaired microglial transition to a disease-associated phenotype and reduced monocyte homing suggests that acquiring the DAM state may depend on signaling from homing monocytes or monocyte-derived macrophages influencing microglial fate. To identify such potential signaling pathways, we performed an *in silico* analysis of cell-cell signaling, focusing on interactions among microglia, monocytes, and macrophages, comparing denervated and sham-operated 5xFAD mice. We applied the MultiNicheNet algorithm, designed to identify differential cell-cell communication networks based on expression levels of ligand-receptor pairs and differential expression of their downstream target genes (43).

Comparing the differential signaling between the sham and denervated 5xFAD mice, we found a complex network of interactions among microglia, monocytes, and macrophages, with significant changes in multiple signaling pathways and their downstream targets across all three cell types (**Fig. 4h-j**). Specifically, following denervation, we observed an overall reduction in microglial signaling, as quantified by the number of differentially expressed ligand-receptor pairs (**Fig. 4h, Extended Data Fig. 3h, i**), with the most pronounced reduction found in microglia-to-monocyte signaling. In contrast, only a small number of signaling pathways were upregulated following denervation.

Notably, we found an increase in signaling from microglia expressing the chemokine *Ccl12*, which is predicted to attract *Ccr2*-expressing monocytes to the brain (also known as monocyte chemotactic protein, MCP-5) (44) (**Fig. 4h**). This elevated signaling reflects a heightened inflammatory condition in response to reduced monocyte homing, consistent with previous reports (9, 10, 15). We identified multiple signaling pathways that were downregulated following splenic denervation in 5xFAD mice. Interestingly, among these down-regulated signaling pathways, we found ligands and receptors associated with the DAM states (**Fig. 4j**). For example, we observed a loss of signaling from monocytes expressing *Apoc2* to microglia expressing *Lpl*. LPL variants have been implicated in genetic risk for AD (45). Another hallmark DAM-associated pathway involved *Trem2*, a well-established genetic risk factor for AD (46–48). We observed a loss of *Apoe*-mediated signaling from monocytes to *Trem2*-expressing microglia following denervation (**Fig. 4j**). Both human and mouse studies have shown that *Trem2* is essential for facilitating the transition of microglia to a disease-associated phenotype in AD (41, 46, 49). Our results are consistent with previous reports showing that *Trem2* deficiency in the AD mouse model at early stages of the disease accelerated cognitive decline (50).

To gain additional insights regarding the downstream target genes of differential signaling following denervation, we constructed a network including the top 150 targets activated by differentially expressed ligands. A notable target gene that was found to be elevated in the sham-operated mice but downregulated following denervation was *Ccl3* (**Fig. 4i**). *Ccl3*, a ligand of the *Ccr5* receptor with autocrine activity on microglia, has been associated with immune cell recruitment and microglial activation (51–53). These alterations in target gene expression, and their regulation by monocytes and microglia, underscore the importance of monocyte-derived signaling in shaping microglial responses in the AD model. Overall, our findings support a model in which monocyte signaling to microglia is critical for the transition to the DAM state, and the reduced abundance of DAM might underlie the observed exacerbation of disease progression at its early stages.

### Splenic denervation impairs monocyte recruitment and exacerbates neuronal loss in a model of acute retinal ganglion cell toxicity

To generalize the role of brain–spleen communication in monocyte recruitment for tissue repair beyond AD, we used a model of acute neuronal injury induced by retinal ganglion cell (RGC) excitotoxicity in WT mice (**Fig. 5a**). This experimental model was chosen based on the previous demonstration that RGC survival is dependent on monocytes (54–56). In addition, due to their location outside the brain, RGCs provide a system that is independent of the contribution of other structures at the other brain’s border, such as the meninges or the choroid plexus. To induce retinal cytotoxicity, we intravitreally injected 80 nmol of L-glutamate to WT mice 2 weeks after splenic denervation (**Fig. 5a**). Retinas were analyzed by flow cytometry 3 days post-injury to assess the abundance of immune cell populations. We found that ethanolization of the splenic nerve significantly reduced monocyte recruitment to the injured retinas (**Fig. 5b, c**), with no effect on T-lymphocyte accumulation (**Fig. 5d**). We repeated the experiment, collecting retinas 7 days after injury to follow RGC survival by immunofluorescence (**Fig. 5e**). We found significantly greater loss of RGCs after retinal injury in the denervated animals compared to sham operated mice (**Fig. 5f**). These findings highlight the importance of intact brain–spleen communication in enabling monocyte homing to sites of neuronal injury and supporting neuronal survival, irrespective of the type of insult.

**Fig. 5:**
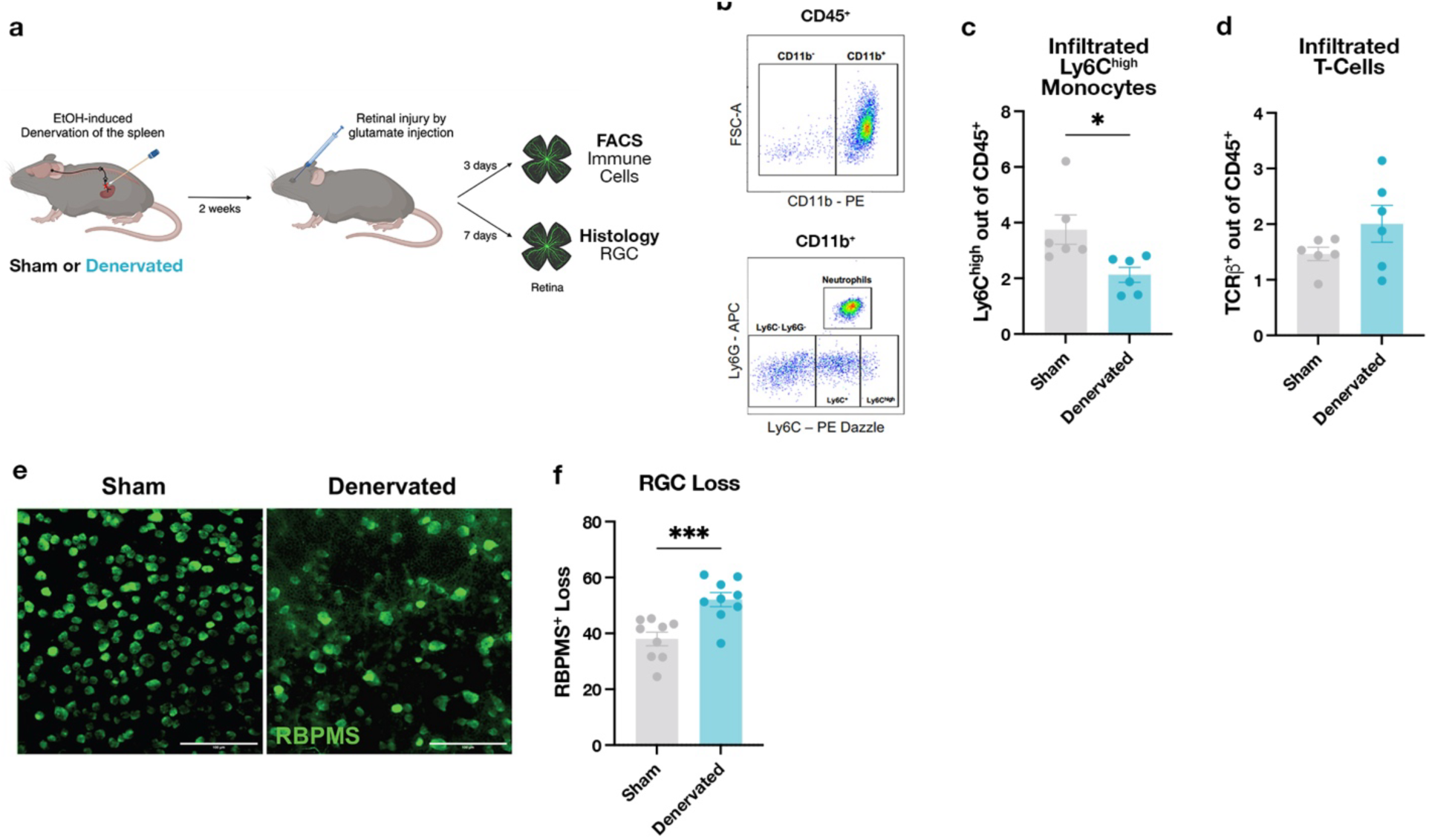
Denervation of the spleen impairs ganglion cell survival following retinal cytotoxicity. **a**, Scheme depicting the experimental design of denervation before the retinal injury. WT mice (n= 6M; n= 6F) were subjected to denervation or sham procedure, and 2 weeks later, received an intravitreal injection of L-glutamate to induce retinal injury. After 3 or 7 days, retinas were analyzed by flow cytometry or immunofluorescence, respectively. **b**, Gating strategy for the detection of myeloid cells in the retina. **c,** Flow cytometry analysis of infiltrated monocytes in the retina of injured animals. **d**, Flow cytometry analysis of T cells in the retina of injured animals. **e**, Representative immunofluorescence of RBPMS in the whole-mount of retinas of injured animals that were previously denervated or sham-operated. Scale bar: 100 μm. **f**, Quantification of RBPMS^+^ retinal ganglion cell (RGC) loss in whole mount retinas of injured animals that were denervated (n= 5M; n= 4F) or sham-operated (n= 5M; n= 4F). Loss of RGC was calculated as the percentage of reduction in the number of RGC compared to uninjured animals. Unpaired t-test. In all panels, data are represented as mean ±SEM. *p < 0.05, ***p < 0.001.

### Increased norepinephrine in the spleen prevents memory loss in 5xFAD mice

Our findings suggested that innervation of the spleen, which mainly relies on sympathetic fibers that release NE, might affect monocyte maturation and migratory capacity. To test this, we increased catecholaminergic tone in the spleen and assessed whether this manipulation could enhance monocyte differentiation, promote monocyte recruitment to the brain, and slow-down cognitive decline in the 5xFAD model of amyloidosis. To this end, we induced overexpression of tyrosine hydroxylase (TH), a rate-limiting enzyme for catecholamine synthesis, in splenic neurons of 7-month-old WT and 5xFAD mice by injecting an AAV2-retrovirus encoding the TH gene under the catecholaminergic neuron–specific PRSx8 promoter (**Fig. 6a**). Control WT and 5xFAD mice received an AAV2-retrovirus encoding GFP driven by the same promoter. Targeted metabolomics confirmed that TH overexpression increased splenic norepinephrine (NE) levels (**Fig. 6b**).

**Fig. 6:**
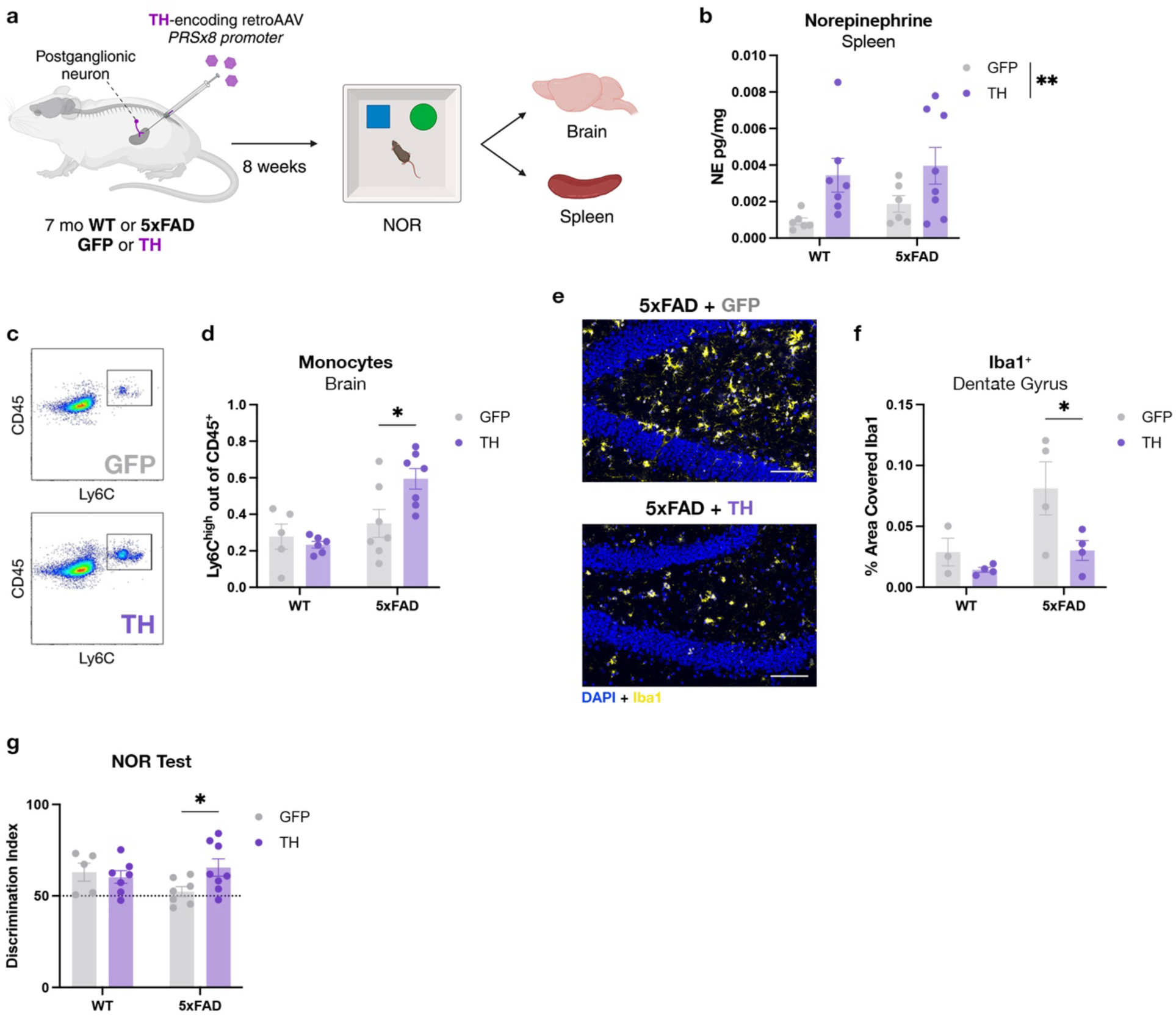
Enhancement of catecholaminergic signaling in the spleen delays symptom onset in the 5xFAD model. **a**, Schematic representation of the experimental setup of tyrosine hydroxylase (TH) overexpression in the 5xFAD model. 7-month-old WT (n= 11M; n= 2F) or 5xFAD (n= 6M; n= 8F) mice were injected in the spleen with a retro-adeno-associated virus (retroAAV) carrying an expression vector for TH or GFP under a catecholaminergic-specific promoter (PRSx8). The NOR test and tissue harvesting were performed 2 months later. **b**, Quantification of norepinephrine levels in the spleen by targeted metabolomics. **c**, Representative flow cytometry gating of monocytes in the brain of 5xFAD animals treated with retroAAV-GFP or retroAAV-TH. **d**, Flow cytometry analysis of infiltrating monocytes in the brains of WT and 5xFAD animals treated with retroAAV-GFP or retroAAV-TH. **e**, Representative images of Iba1 staining in the dentate gyrus (DG) of 5xFAD mice overexpressing GFP or TH in the spleen. Scale bar: 100 μm. **f**, Iba1 quantification by immunofluorescence in the DG of WT and 5xFAD overexpressing GFP or TH in the spleen. **g**, Discrimination index in the novel object recognition (NOR) test, calculated as the percentage of time spent exploring the novel object over the total exploration time of the two objects, by WT and 5xFAD mice, 2 months after TH or control virus injection. Two-way ANOVA followed by Sidak’s multiple comparison test. In all panels, data are represented as mean ±SEM. *p < 0.05, **p < 0.01.

We next assessed the effect of increased NE on splenic hematopoiesis. Flow cytometry quantification of HSPC subsets revealed no change in total LSK numbers (**Extended Data Fig. 5a, b)**. However, multipotent progenitors with lymphoid and myeloid potential (MPP3/4) were significantly increased in TH-overexpressing 5xFAD mice compared with GFP controls (**Extended Data Fig. 5c**), indicating that TH overexpression enhanced splenic extramedullary hematopoiesis.

Next, we used flow cytometry to assess the effect of TH overexpression in the neurons that innervate the spleen on monocyte recruitment to the brain. We observed increased monocyte infiltration in the brains of TH-5xFAD mice, but not in TH-WT mice (**Fig. 6c, d**). This result supports the notion that spleen catecholaminergic signaling plays a role in regulating myeloid cell recruitment to the CNS upon need. Histological analysis revealed a decrease in the Iba1 signal in the DG (**Fig. 6e, f**), along with a trend toward higher synaptophysin levels in the CA3 region of TH-5xFAD mice compared to GFP-5xFAD controls (**Extended Data Fig. 5d, e**), indicating synaptic rescue in the diseased brain when monocyte recruitment was augmented. Finally, we examined whether increased NE signaling in the spleen could attenuate cognitive decline. In the NOR test, 9-month-old 5xFAD mice with splenic TH overexpression showed significantly better recognition memory 2 months after treatment than GFP controls (**Fig. 6g**). Overall, these findings suggest that boosting catecholaminergic activity in the spleen can delay disease symptoms in a mouse model of neurodegeneration.

## Discussion

In this study, we show that disruption of brain–immune communication contributes to AD progression. Mechanistically, this pathway depends, at least in part, on peripheral monocyte recruitment and on monocyte–microglia interactions in the brain. We first showed that brain–spleen communication is impaired in AD mice at a stage when cognitive deficits are present. When we experimentally disrupted this pathway at the presymptomatic stage, it was sufficient to accelerate cognitive decline. As a corollary, enhancing NE signaling in the spleen attenuated disease manifestations.

The brain’s ability to regulate the immune response through various pathways has been a subject of extensive research (57–60), with significant implications for neurodegenerative diseases. AD has been increasingly linked to systemic immune dysfunction and local brain inflammation (61, 62). Yet, the precise mechanisms by which the brain influences peripheral immune responses, and the reciprocal effects of the immune response on the brain, remain only partially understood. Previous studies demonstrated that the brain can sense and signal to the spleen to mobilize immune cells (26, 63, 64). By selectively disrupting the brain–spleen axis without directly manipulating either the brain or immune cells, we investigated how this cross-talk influences both systems, highlighting their interdependence.

It is clear that both descending efferent signals and ascending sensory inputs contribute to this interaction, and both may have been affected by our surgical procedure. Moreover, monocytes can reach the brain through several routes, including skull microchannels that connect bone marrow to the meninges (65–67). Nevertheless, consistent observations in both WT mice after retinal injury and in 5xFAD mice after splenic denervation reinforce the spleen’s role as a critical reservoir for monocytes, determining their capacity to migrate to the brain when needed. While in the 5xFAD model, our observations highlight the spleen as an important site of extramedullary hematopoiesis, modulated by noradrenergic tone, we cannot exclude potential contributions from other monocyte reservoirs and sources, such as the skull or peripheral bone marrow.

Cognitive and functional deficits in AD models result from several factors that go awry in the brain, including the accumulation of protein aggregates, altered glial states, and impaired neuronal function. At the early stages of disease, while amyloid plaques accumulate, there are no signs of cognitive deficit (68). We found that denervation of the spleen at this early stage accelerated cognitive decline, yet this was not associated with increased plaque burden, but rather with significant changes in the activation state of microglial cells and in neuronal synaptic integrity. Our findings are aligned with previous reports demonstrating microglial control of synaptic integrity (69–71), which was found here to be negatively affected following spleen denervation. Furthermore, our findings revealed an intimate interaction between monocytes, microglia, and macrophages, involving genes required for microglial acquisition of the DAM state, a differentiation state previously shown to be beneficial at the early stages of the disease (46, 72–74). Moreover, these results align with prior observations that boosting systemic immunity by blocking the immune checkpoint PD-L1/PD-1 pathway improved outcomes in multiple AD mouse models through a monocyte-dependent mechanism and was accompanied by increased microglial representation of a disease-associated (DAM) state (14). Our results highlight a potential novel mechanism through which peripheral myeloid cells are recruited to the CNS under pathological conditions, beyond their anti-inflammatory and phagocytic activities^17^.

Considering the significance of monocytes and monocyte-derived macrophages in regulating the disease process, it is noteworthy that there is no noticeable increase in these protective cells during disease progression. This observation is particularly perplexing in light of the microchannels directly connecting the skull’s bone marrow to the meninges (65, 75, 76). A disruption of the brain-spleen connection, combined with the increased presence of soluble factors that inhibit monocyte differentiation (77), occurring as pathology advances, may partly explain this detrimental limitation.

Building on our findings, we propose that monocytes support resident CNS cells in preserving brain function during a clinically silent phase of AD that can last for decades. As the pathology progresses and affects brain regions critical for brain–immune communication, this signaling becomes disrupted, limiting the spleen’s ability to supply protective monocytes that interact with microglia and shape their response. Taken together, our results support the hypothesis that brain pathology in AD extends beyond the CNS to influence other organs and systems, including the immune system, and that reciprocal immune dysfunction, in turn, has profound consequences for disease progression.

## Methods

### Mice

All experiments described here complied with the regulations formulated by the Institutional Animal Care and Use Committee (IACUC) of the Weizmann Institute of Science. Female and male mice were bred and maintained by the animal breeding center of the Weizmann Institute of Science. For all experiments, heterozygous 5xFAD transgenic mice (line Tg6799, The Jackson Laboratory) on a C57/BL6-SJL background and wild-type (WT) littermates were used. Genotyping was performed by PCR analysis of ear clip DNA, as previously described. Mice subjected to the various manipulations were mixed within cages. On the day of sacrifice, mice were anesthetized and transcardially perfused with ice-cold PBS. For histology, after dissection, the brain was post-fixed in 4% paraformaldehyde (PFA) in PBS. For immune profiling of brain immune cells by Flow Cytometry or scRNAseq, the brains were excised excluding the medulla, without dissecting meninges and choroid plexus, and manually chopped (0.5–1.0 mm^2^ in size), before enzymatic dissociation using 0.4mg/ml Collagenase D, 2mM HEPES (BI Industries), 10μg/ml DNase (Sigma), and 2% FCS for 20 min at 37°C. For density gradient separation, the pellet was resuspended with 37% Percoll in PBS and centrifuged at 800g for 20 min at 4°C with no brake; the supernatant was then discarded. Cells were suspended in ice-cold sorting buffer (PBS supplemented with 2 mM EDTA and 2% FCS) supplemented with anti-mouse CD16/32 (1:100, no. 101302, BioLegend) to block Fc receptors before labeling with fluorescent antibodies against cell-surface epitopes (Supplementary Table 1). For scRNA sequencing, samples were gated for CD45^+^ CD11b^+^ after exclusion of debris and doublets, and sorted using an ARIA-III instrument (BD Biosciences), and analyzed with BD FACSDiva (BD Biosciences) software. Isolated single cells were sorted into 1.5ml Eppendorf tubes containing PBS enriched with 10% ultra-pure BSA and then immediately subjected to library preparation for scRNA sequencing. For immune profiling of the spleen either by CyTOF or scRNA sequencing, the organ was mashed with a syringe plunger against a 70 mm strainer, and treated with RBC Lysis Buffer (BioLegend) to remove erythrocytes before labeling with antibodies.

For the experiments employing glutamate excitotoxic-induced damage of the retina, WT mice were anesthetized, and local anesthesia (Localin; Dr. Fischer) was applied directly to the eye. Mice were injected intravitreally with a total volume of 1 μl saline containing 80 nmol l-glutamate (Sigma-Aldrich), as previously described (78). At the indicated timepoints post-injury, mice were anesthetized, perfused intracardially with PBS, and their retinas removed by dissection. For Flow-cytometry analysis, retinas were digested with using 0.4mg/ml Collagenase D, 2mM HEPES (BI Industries), 10μg/ml DNase (Sigma), and 2% FCS for 10 min at 37°C, and then filtered through a 70μm strainer.

### In vivo anti-CCR2 Injections

Anti-CCR2 (clone MC-21) was administered at 400 μg per injection in 200 μl PBS, every 4 days for 4 injections (last injection 3 days before sacrifice).

### Histology and immunohistochemistry

For brain sections, paraffin-embedded tissue was sectioned at a thickness of 6 μm. One slide per animal was used for staining, each containing 6 equally spaced sections. Antibody staining was performed as previously described(79), except that all primary antibodies were incubated overnight at RT, followed by another overnight at 4°C. For Aβ staining, the Mouse On Mouse detection kit (Vector labs) was used according to the manufacturer’s instructions. The following primary antibodies were used: mouse anti-human Aβ (1:150; Covance); chicken anti-GFAP (1:150; Abcam), anti-Synaptophysin (1:200, Abcam), anti-Iba1 (1:150, Wako). For secondary antibodies, Cy2/Cy3-conjugated anti-mouse/chicken (1:150; Jackson Immunoresearch) were used. Counterstaining was performed using 4’,6-diamidino-2-phenylindole (1:5000; Biolegend).

For imaging of the retina, after cornea and lens removal in PBS, eyecups were fixed for 1 hour (4% PFA at 4°C) and then hemisected to obtain wholemount retinas. Retinal wholemounts were washed in PBS, blocked with 3% BSA in 0.3% Triton X-100 in PBS for 2 hours at room temperature and then incubated for 3 days in primary antibody solution (1% BSA dissolved in 0.1% Triton X-100 in PBS; primary antibody: rabbit polyclonal anti-RBPMS antibody, 1:100 (PhosphoSolutions, Cat. No. 1830)) at 4°C on a shaker. The next day, retinas were washed in PBS, and immersed overnight in a secondary antibody solution. The tissues were stained with DAPI to identify nuclei and mounted onto Superfrost/Plus Microscope Slides, and covered with a coverslip, using Aqua-Poly/Mount (Polysciences) mounting medium. Retinal wholemounts were digitally scanned using Olympus UPlanSApo 20x/0.75 NA objectives on an Olympus BX61VS slide scanner (Olympus Corporation, Tokyo, Japan). Further image processing for wholemount retinas was performed with Fiji.

### Targeted Metabolomics

For the detection of Norepinephrine, Epinephrine, Dopamine, Serotonin and Acetylcholine in both plasma and spleen homogenates, targeted metabolomics was used. For plasma, blood was collected in tubes containing 2 μL heparin, briefly vortexed, and then centrifuged at 3,000 g for 15 min at 4 °C. Supernatant plasma was aliquoted and stored at ULT. The spleens were weighed and then disrupted using scissors. For each milligram of spleen, 5 µL of PBS1x with 1:1000 of Protease Inhibitor Cocktail (Sigma, P8340) was added, and the tissue was thoroughly resuspended. The disrupted spleen was transferred to a tissue homogenizer. After homogenization, the homogenate was centrifuged at 10,000g for 10 minutes, and supernatant was collected.

LC-MS/MS analysis was performed according to a previously described protocol (80) with slight modifications as detailed below. A 10-µl aliquot of each sample and 10 µl of internal standard (IS; norepinephrine-d6, dopamine-d4, and serotonin-d4 from C.D.N. Isotopes, 400ng/ml each) were added to 70 µL of borate buffer (200 mM, pH 8.8) at 25°C, mixed, and then 20 µl of Aqc reagent (10 mM dissolved in 100% ACN) was added and immediately mixed. The samples were heated at 55°C for 10 minutes, then filtered into nano filter vials (0.2-µm PES, Thomson) for analysis.

The LC-MS/MS instrument consisted of an Acquity I-class UPLC system and Xevo TQ-S triple quadrupole mass spectrometer (Waters). Chromatographic separation was done on a 100 x 2.1-mm i.d., 1.8-µm UPLC HSS T3 column (Waters Acquity) with mobile phases A (0.1% formic acid in water) and B (0.1% formic acid in acetonitrile) at a flow rate of 0.6 ml/min and column temperature 45°C. The gradient used was as follows: 0.5 min at 4% B; linear increase to 10% B over 2 min, increase to 28% B over 2.5 min, then washing by increase to 95% B in 0.1 min, followed by 2 min hold at 95% B, back to 4% B over 0.1 min, and equilibration at 4% B for 2.8 minutes. Samples kept at 4°C were automatically injected in a volume of 1 µl.

Mass detection was carried out using electrospray ionization in positive mode. Argon was used as the collision gas with flow 0.10 ml/min. The capillary voltage was set to 3.0 kV, source temperature - 150°C, desolvation temperature - 650°C, desolvation gas flow - 800 L/min, cone voltage 20V, collision energy 25eV. Analytes were detected by monitoring of fragment ion 171 m/z produced from corresponding precursor ions: 354 [epinephrine + Aqc + H]+, 347 [serotonin + Aqc + H]+, 340 [norepinephrine + Aqc + H]+, and 324 [dopamine + Aqc + H]+ m/z; for IS: 351 [serotonin-d4 + Aqc + H]+, 346 [norepinephrine-d6 + Aqc + H]+, and 328 [dopamine-d + Aqc + H]+ m/z. Standard curves of the neurotransmitter mix at a range of 0.001-10 uM were used. MassLynx and TargetLynx software (v.4.2, Waters) were used to acquire and analyze the data.

### Cognitive tests

RAWM was used to test hippocampal-dependent spatial learning, following the protocol of Alamed and colleagues (81) with some modifications. Briefly, six stainless-steel inserts were placed in a plastic pool, forming six open and connected arms. A hidden platform was placed at the end of a ‘goal arm’ (arm 6; Extended Data Fig. 3a). Milk powder was used to render the water opaque, and water was maintained at a temperature of 23 ± 1 °C. On day 1, the training phase, mice were subjected to 15 trials. In each trial, mice were allowed 60 s to find the platform. Mice that failed to find the platform were placed on it by the experimenter. The inter-trial interval was, on average, 20 min. Trials alternated between a visible and hidden platform. However, from trial 12 and throughout the second day, the platform was hidden. Spatial learning and memory were measured by an investigator who was blinded to the treatment of the mice, and who recorded the number of arm entry errors (error was defined as entrance to an incorrect arm, or failure to enter any arm within 15 s) as well as the escape latency of the mice on each trial. The 30 trials were grouped into three trial bins—five bins each on days 1 and 2. Data were analyzed by a team member who did not perform the experiment. Animals that showed floating behavior or motor difficulties and were unable to execute the task were excluded from the analysis.

The NOR protocol was modified from Bevins and Besheer (82) and used a 41.5 × 41.5 cm^2^ gray arena. The experiment spanned 2 days and included three trials: (1) a habituation trial—a 20-min session in the empty apparatus (day 1); (2) a familiarization trial—a 10-min session presenting two identical objects located 15 cm apart (day 2); and (3) a test trial. For the test trial, following a 1-h training-to-testing interval, each mouse was returned to the apparatus for a 6-min session in which one of the objects was replaced by a novel one. Mouse behavior was recorded and analyzed by an investigator who was blinded to the treatment group. Novel object preference was defined as ‘discrimination ratio’: time (s) spent interacting with the novel object/(time spent with the familiar object + time spent with the novel object).

### Splenic Denervation

Mice were anesthetized with isoflurane. Splenic denervation was performed following the protocol of Zhang with minor adjustments (26). The peritoneal cavity was accessed through an abdominal incision on the left flank. Forceps were used to isolate the spleen away from the peritoneal cavity so that the three main supplying vasculature trees were clearly exposed. Absolute ethanol was repeatedly applied with arrow-shaped cotton tips to those vasculature trees for 7 seconds each time, at 5-second intervals, and about seven times in total to deplete the splenic nerve fibers that run along them. Care was taken to avoid excessive ethanol dripping from the cotton tip and to avoid causing visible vessel spasms, which could lead to permanent damage to blood vessels and splenic necrosis.

For sham-operated mice, the entire surgical procedure was identical except that saline (pH 7.4) instead of absolute ethanol was repeatedly applied. Mice were sutured using absorbable and silk lines and allowed to recover at a controlled temperature under a heating lamp. Mice recovered within minutes from anesthesia withdrawal and their well-being was monitored daily for the 3 days following surgery or until complete recovery. After denervation surgery, animals were allowed to recover for 4 weeks before cognitive tests.

### Viral constructs and injection

The Pseudorabies virus carrying RFP (PRV_mRFP_VP26 also known as PRV 765) was previously described(83). The virus was used for retrograde neuronal labeling at a titer of 1.5*10^8 PFU/ml.

For overexpression of Tyrosine Hydroxylase in the splenic nerve, the rAAVrg-PRSx8-TH virus was used (3.58*10^10 PFU/ml), and rAAVrg-PRSx8-eGFP (3.75*10^9 PFU/ml) was used as a control.

The PRSx8 promoter was amplified from pAAV-PRSx8-eGFP plasmid (Addgene #192589)by PCR using Fw-tcggcaattgaaccggtTTTAATTAAAAACGCGTATT and Rv-agaGTCGACggtacctcacgacacTCTAGAGTCGGCTGGGGTGAGCTCTCTGGTCCCACCTGGCCA containing an overhang of SalI restriction sites (84). Afterwards, the promoter of pAAV-Ef1α-DIO-TH-p2A-mRFP was exchanged with the PRSx8 (85) to obtain pAAV-PRSx8-TH. AAV generation:

To produce rAAV2-retro, a triple co-transfection procedure was used to introduce a rAAV vector plasmids together with rAAV2-retro, helper plasmid carrying AAV *rep* and *cap* genes (rAAV2-retro helper was a gift from Alla Karpova and David Schaffer (Addgene plasmid #81070; http://n2t.net/addgene:81070; RRID:Addgene_81070)) and pXX6-80, Ad helper plasmid, at a 1:1:1 molar ratio(86).

Briefly, HEK293T cells were transfected using poly- ethylenimine (PEI) (linear; molecular weight [MW], 25,000) (Poly- sciences, Inc., Warrington, PA), and medium was replaced at 18 h post- transfection. Cells were harvested at 72 h post-transfection, subjected to three rounds of freeze-thawing, and then digested with 100 U/ml Benzonase (EMD Millipore, Billerica, MA) at 37°C for 1 h. Viral vectors were purified by iodixanol (Serumwerk Bernburg AG, Germany) gradient ultracentrifugation (according to Zolotukhin et al, Gene Therapy (1999) 6, 973–985) followed by further concentration using Amicon ultra-15 100K (100,000-molecular-weight cutoff, Merck Millipore, Ireland) and washed with phosphate-buffered saline (PBS -/-). Final concentration of rAAV2-retro particles was 7.56E+10 ng per microliter (PRSx8_eGFP), and 3.58E+10 ng per microliter (PRSx8_ TH).

**For intrasplenic injection**, the spleen was exposed as described above, and 4µl of viral suspension was injected, 2µl in the upper pole and 2µl in the lower pole, using a 33G gauge needle.

**The locus coeruleus was targeted** using stereotactic injection. Anesthetized mice were fixed with an auxiliary ear bar (EB-6, Narishige Scientific) in a stereotactic frame and positioned on a heating blanket to maintain body temperature. Eyes were covered with an eye ointment to avoid drying. Subsequently, the scalp was opened (∼1 cm incision) and the skull was cleaned with NaCl. Bregma and lambda were identified and stereotactically aligned. A small craniotomy was then performed above the locus coeruleus (AP: –5.4; ML: ±01.1; DV: –3.6 mm relative to bregma) using a dental drill that was constantly kept moist. The viral suspension was drawn into a microsyringe (RWD) and 0.3 μl volumes were bilaterally injected into the LC with an injection speed of ∼100 nl/min using an injection pump. The needle was kept in place for at least 2 minutes before injection and 10 min after finishing the injection to avoid undesired viral spread and was then slowly withdrawn. The skin was closed with simple interrupted sutures and disinfected with iodine. Mice received analgesia and were placed in a cage on a heating pad for recovery.

### CyTOF

For the characterization of immune cells in the spleen and the brain, dedicated CyTOF panels were used and in-house conjugations were performed using the MIBItag Conjugation Kit (IONpath) where needed.

To characterize brain immune cells, after the single-cell extraction, samples were incubated for 5 min with anti-mouse CD16/32 (1:100, no. 101302, BioLegend) and then barcoded using anti-CD45 antibodies labelled with different isotopes, as previously described(87). Cells were washed twice with MaxPar Cell Staining Buffer (Fluidigm) followed by incubation with the extracellular antibodies (30min, RT). Then, the samples were fixed and resuspended with 4% Formaldehyde (Pierce) for 10 min, washed, and kept on ice for 10 min. They were then permeabilized using ice-cold 90% methanol for 15 min on ice, and stained for intracellular markers in the presence of 1% donkey serum for 30 min. Finally, the samples were washed twice and kept in 4% Formaldehyde (Pierce) with Iridium (125 nM) at 4°C overnight. On the day of analysis, samples were washed twice with MaxPar Cell Staining Buffer (Fluidigm), then twice with MaxPar water (Fluidigm), resuspended at a concentration of 300K cells/ml in 1:10 EQ Four Element Calibration Beads (Fluidigm) in MaxPar water, and filtered via a 35 um mesh before acquisition using a CyTOF 2 upgraded to Helios system (Fluidigm). A detailed list of antibodies used is supplied in Supplementary Table 2. After normalization, gating of live single cells and debarcoding, the analysis was performed using Cytof Workflow (88). For the analysis of splenocytes, only surface staining was used.

### Pre-processing of scRNA-Seq data

Brain and spleen single-cell RNA libraries were prepared using 10x Chromium Next Gem Single cell 3’ reagent kit (version 3.1) according to the manufacturer’s instructions, and quantified using High Sensitivity dDNA Reagents on the Qubit System, and High Sensitivity D1000 Reagents (5067- 5582) on the 2200 TapeStation system (Agilent). The libraries were sequenced on Illumina’s NovaSeq 6000 platform.

Spleen samples were demultiplexed using mkfastq. Cell assignment, alignment to the mm10 mouse genome, and count matrix generation were done with CellRanger multi command (version 6.0.1). For brain samples, cell assignment, alignment to the mm10 mouse genome, and count matrix generation were performed with the CellRanger count command (version 6.0.1).

To account for technical artifacts in the data, specifically correcting gene counts shifted due to ambient RNA, the CellBender(89) (version 2) program was run on each sample, to remove counts due to ambient RNA molecules and random barcode swapping from (raw) UMI-based single cell or nucleus RNA-Seq count matrices, and to determine the cell barcodes that are valid cell libraries, excluding empty droplets and low-quality libraries. CellBender output was used as input for downstream analysis.

For every cell, the number of genes for which at least one read was mapped was quantified, and then cells with fewer than 200 detected genes were excluded. We removed mitochondrial genes, and non-coding genes with no canonical names. The SCTransform(90, 91) function from the Seurat package (version 5.0.1), was used for normalization and variance stabilization of single-cell RNA-seq data using regularized negative binomial regression.

### Dimensionality reduction, clustering, and visualization

Clustering and downstream analysis were done using the Seurat package version 5.0.1. The normalized RNA count matrix was then used for dimensionality reduction, visualization, and clustering. Dimensionality reduction was done with principal component analysis (PCA, using RunPCA function in Seurat). After PCA, significant principal components (PCs) were identified using an elbow plot of the distribution of standard deviation of each PC (ElbowPlot in Seurat). In both analysis of brain and spleen datasets, 20 PCs were used for the analysis of all cells, 10 PCs for microglia, monocyte and macrophage analysis, and 10 for microglia-only analysis. Within the top PC space, transcriptionally similar cells were clustered together using a graph-based clustering approach. First, a k-nearest neighbor (k-NN) graph was constructed based on the Euclidean distance. For any two cells, edge weights were refined by the shared overlap of the local neighborhoods using Jaccard similarity (FindNeighbors method Seurat, with k = 30). Next, cells were clustered using the Louvain algorithm(92) which iteratively grouped nuclei and located communities in the input k-NN graph (FindClusters function in Seurat, with resolution 0.8 for all brain and spleen cells, resolution 0.5 for microglia, monocyte and macrophage, and resolution 0.6 for microglia only cells). For visualization, the dataset was embedded in 2 dimensions by UMAP, using the same top principal components as input to the algorithm (using the RunUMAP function in Seurat).

For sub-clustering analysis within cell types, each cell type (i.e. microglia, monocytes, and macrophages in the brain tissue) was subsetted from the main dataset for a high-resolution analysis (using the PCs and resolution mentioned above). Cells were clustered at high resolution, and clusters were then annotated and merged based on known marker gene expression and RNA profile similarity.

### Annotation of clusters to cell types

Identification of cell types was done using the Bioconductor package SingleR (93), which performs unbiased cell type recognition from single-cell RNA sequencing data, run with ImmGenData from the Immunologic Genome Project(94). Cell types were annotated according to the classification, and further validated using the expression of known marker genes (3).

### Differential expression analysis

Differentially expressed genes (DEGs) between the two conditions within the monocytes (infiltrating and patrolling clusters) were identified using the Wilcox test implementation in Seurat’s FindMarkers function (assay set to integrated). P-values were adjusted for multiple hypothesis testing using Benjamini-Hochberg’s correction (FDR). Significance was set to an FDR-adjusted P-value threshold of 0.05 and a log fold change threshold of 0.25. Volcano plots were generated using ggplot2.

### Cell-cell communication

The MultiNicheNet package (version 1.0.3)(43) was used to determine interactions between cell clusters based on annotated interactions of ligands, receptors and the downstream genes within signaling pathways, and the level of expression of ligand and receptors within our scRNA-seq dataset. Each condition was compared to the two other conditions to determine specific cell-cell communication in the condition of interest. The default MultiNicheNet pipeline was used, except for the parameter minimum logFC threshold that was set to 0.1. Microglia, monocytes, and macrophages were used for this analysis as both sender and receiver cells. First, cell-cell communication between all clusters was determined, and the top 30 predictions per condition were selected for visualization in a Circos plot. Next, the prioritization table was filtered to attain the top 10% scores (using a prioritization score threshold larger than 0.716), and all the predictions including DAM marker genes^20^ were selected for visualization in a Circos plot. To visualize the total interactions, Sankey plots were used, prepared using the Sankey Network function of the networkD3 package version 0.4, showing the number of edges between ligand-expressing and receptor-expressing cell types. Two Sankey plots were combined to show the differential graphs between the two conditions of sham vs. denervated 5xFAD mice. Finally, we generated a systems view plot of the intercellular feedback and cascade processes than can be occurring between the different cell populations involved. Links were drawn between ligands of sender cell types and their ligand/receptor-annotated target genes in receiver cell types.

---

**Extended Data Fig. 1:**
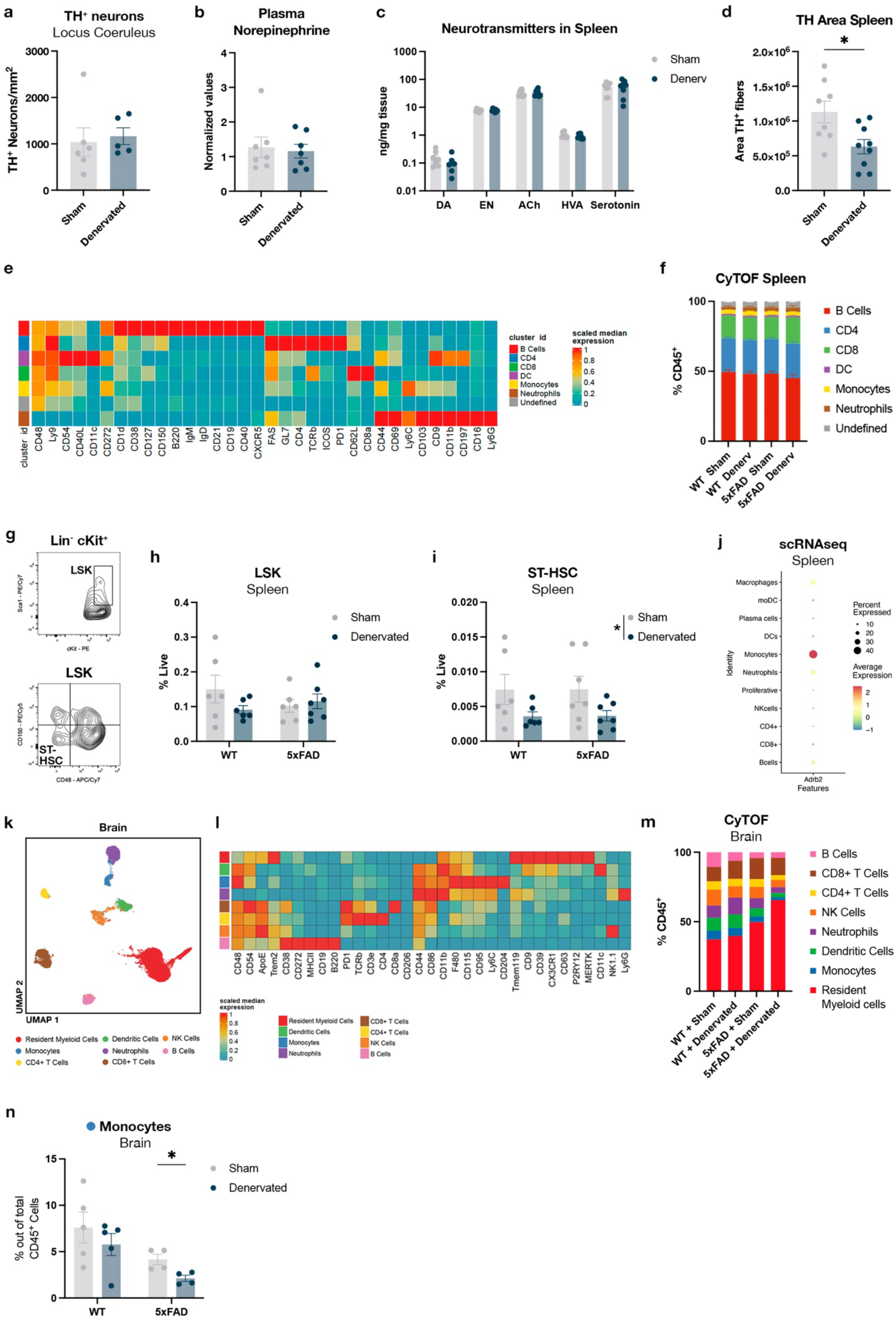
Spleen denervation in the 5xFAD mouse model causes alterations in extramedullary hematopoiesis and increases monocyte homing to the brain. **a**, Quantification of TH^+^ neurons in the locus coeruleus of denervated animals compared to sham-operated mice. **b**, Levels of systemic circulating norepinephrine in the plasma of sham-operated and denervated mice. **c**, Levels of neurotransmitters detected in the spleen of denervated animals compared to sham operated mice. DA, dopamine; EN, epinephrine; ACh, acetylcholine; HVA, homovanillic acid. **d**, Quantification of tyrosine hydroxylase (TH) by immunofluorescence in the spleen of sham and denervated animals. **e**, Heatmap with scaled marker expression values detected by mass cytometry (CyTOF) in the spleen of sham-operated or denervated WT or 5xFAD animals. **f**, Proportions of different immune cell subsets detected by CyTOF in the spleen of sham-operated or denervated WT or 5xFAD animals. **g**, Flow cytometry gating strategy of HSPC subsets in the spleen of 5xFAD mice. **h**, Abundance of Lin^-^Sca1^+^cKit^+^ (LSK) hematopoietic progenitors in the spleen of sham-operated or denervated WT or 5xFAD mice. **i**, Short-term hematopoietic stem cells (ST-HSC) in the spleen in WT or 5xFAD animals after denervation, compared to sham-operated animals. **j**, Dot-plot showing the scaled expression of the *Adrb2* gene in the different cell types found in the spleen by single-cell RNAseq. **k**, Dimensional reduction by UMAP of mass cytometry (CyTOF) data of immune cells in the brain of sham or denervated WT and 5xFAD mice. **l**, Heatmap showing the scaled signal of markers studied by CyTOF to annotate clusters of immune cells. **m**, Proportion of the different immune cell types detected in the brains of WT and 5xFAD mice that were denervated or sham-operated. **n**, Proportion of infiltrating monocytes in the brain as determined by CyTOF. Unpaired t-test or two-way ANOVA followed by Sidak’s multiple comparisons test. In all panels, data are represented as mean ±SEM. *p < 0.05.

**Extended Data Fig. 2:**
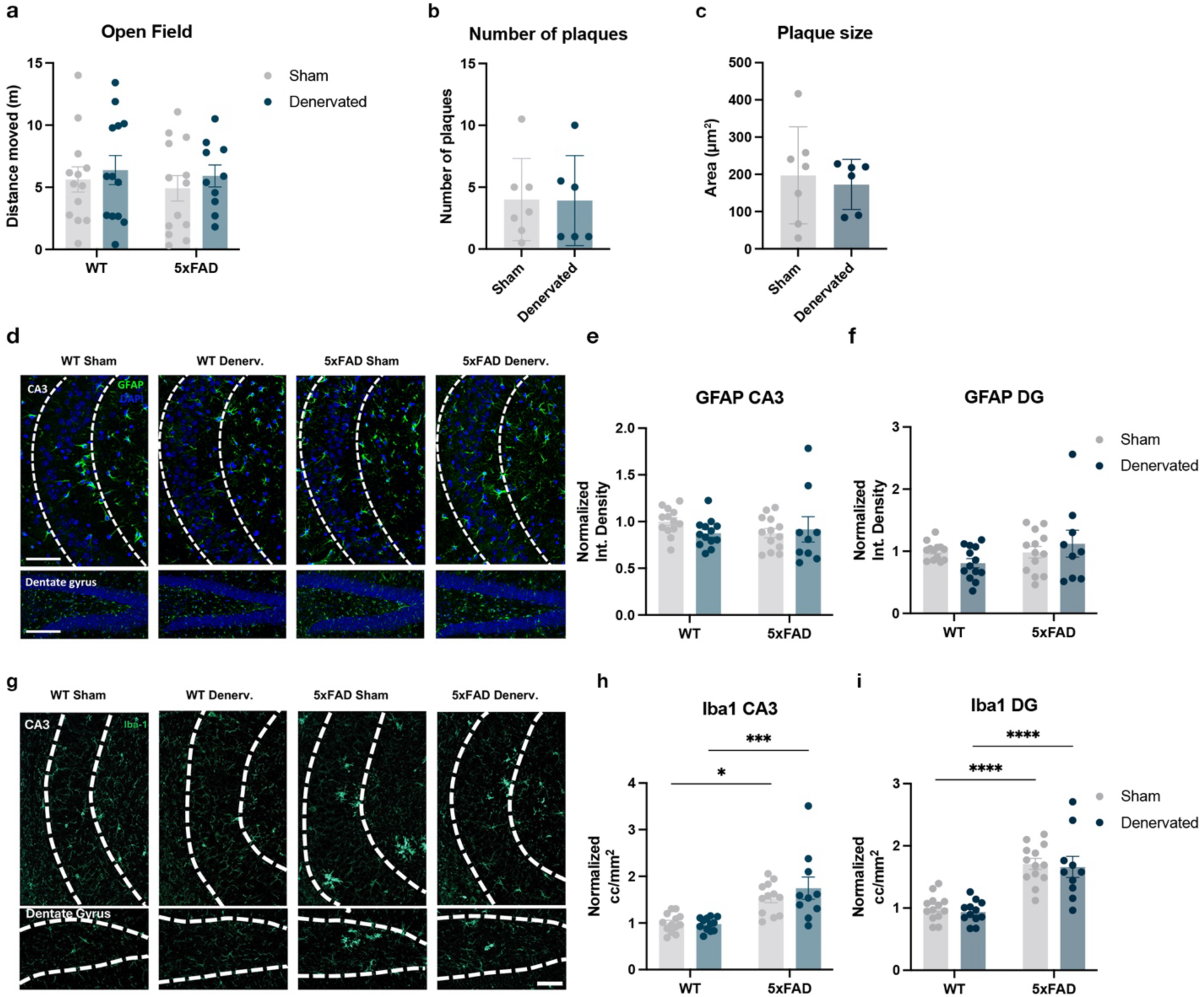
Denervation of the spleen does not affect amyloid deposition, astrogliosis or microgliosis in the 5xFAD model. **a**, Distance moved during 20 minutes in the open field test. **b**, Number of plaques in the hippocampus of sham-operated or denervated 5xFAD animals. **c**, Plaque size in the hippocampus of sham-operated or denervated 5xFAD animals. **d**, Representative images of GFAP immunolabeling in the CA3 and the dentate gyrus in the hippocampus of sham-operated or denervated WT or 5xFAD mice. Scale bar: 100 μm. **e**, Quantification of the GFAP intensity in the CA3 of sham-operated or denervated WT and 5xFAD animals. **f**, Quantification of the GFAP intensity in the DG of sham-operated or denervated WT and 5xFAD animals. **g**, Representative images of Iba1 immunostaining in the CA3 and the dentate gyrus in the hippocampus of sham-operated or denervated WT and 5xFAD mice. Scale bar: 100 μm. **h**, Quantification of the normalized Iba1 counts in the CA3 of sham-operated or denervated WT or 5xFAD animals. **i**, Quantification of the normalized Iba1 counts in the DG of sham-operated or denervated WT or 5xFAD animals. Unpaired t-test or two-way ANOVA followed by Sidak’s multiple comparisons test. In all panels, data are represented as mean ±SEM. *p < 0.05, ***p < 0.001.

**Extended Data Fig. 3:**
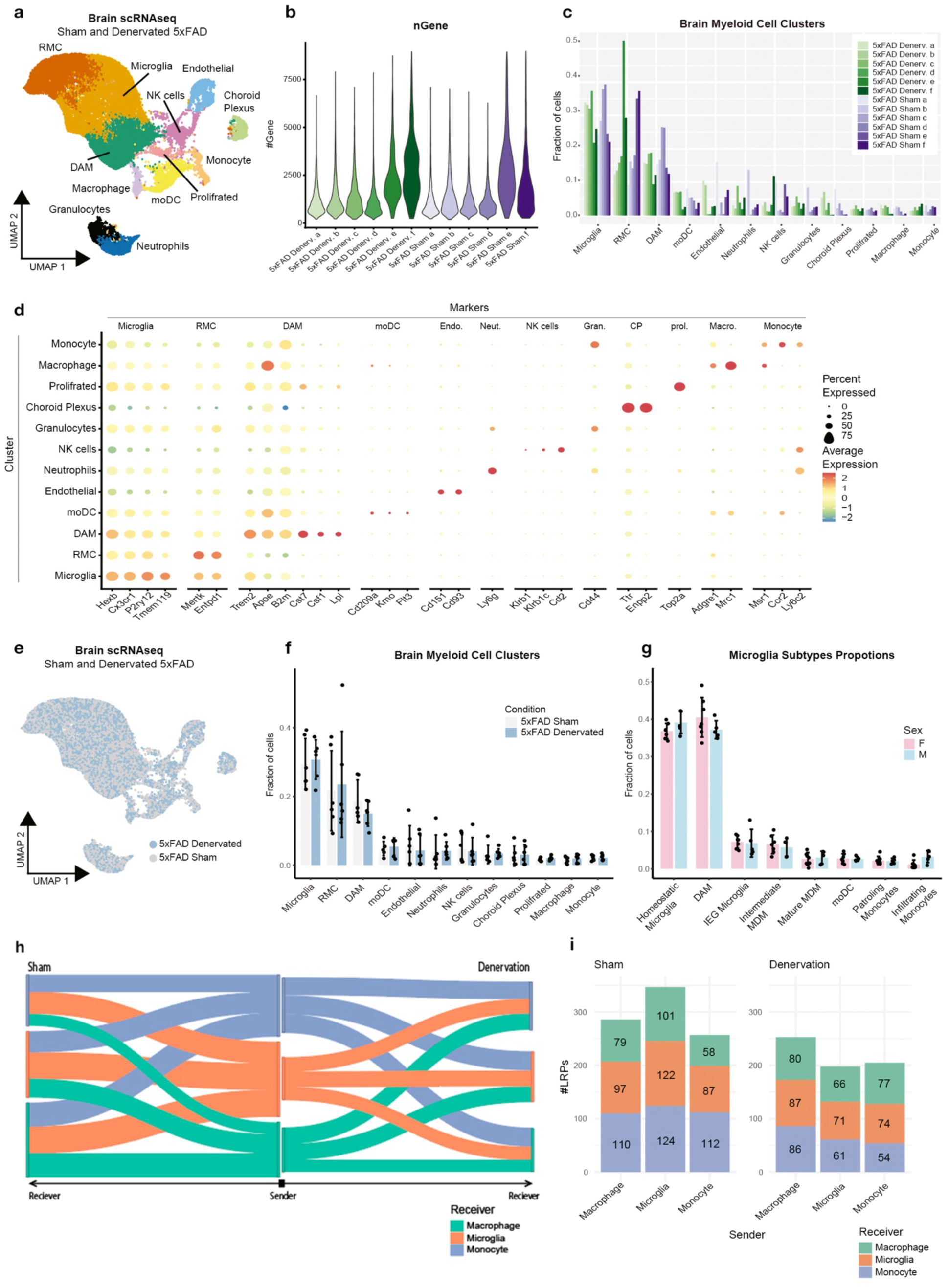
Denervation of the spleen remodels monocyte-microglia communication in the brains of 5xFAD mice. **a**, UMAP embedding of 34,198 single-cell RNA profiles from whole brain- myeloid cells from sham-operated or denervated 5-month-old 5XFAD mice. **b**, Number of genes across samples. Violin plot showing the distribution of the number of transcripts (unique UMIs) detected per mouse. **c**, Bar plot of the proportions of all cell types recognized in the myeloid cell-enriched dataset across samples. **d**, Marker genes used for annotation of the clusters. **e**, UMAP embedding of 34,198 single-cell RNA profiles from whole brain- myeloid cells from sham-operated or denervated 5-month-old 5XFAD mice, color-coded by treatment. **f**, Bar plot of the proportions of brain-myeloid cell types across the two conditions: sham-operated and denervated. **g**, Bar plot of the proportions of brain-myeloid cell types across sex: Male and Female. **h**, Number of statistically significant LRPs between pairs of cell types (color-coded) across experimental groups out of top 25% ranked LRPs of the MultiNicheNet analysis (score>0.653). **i**, Differential ligand-receptor pairs (LRPs) across experimental groups out of top-ranked LRPs of the MultiNicheNet analysis (score>0. 653).

**Extended Data Fig. 4:**
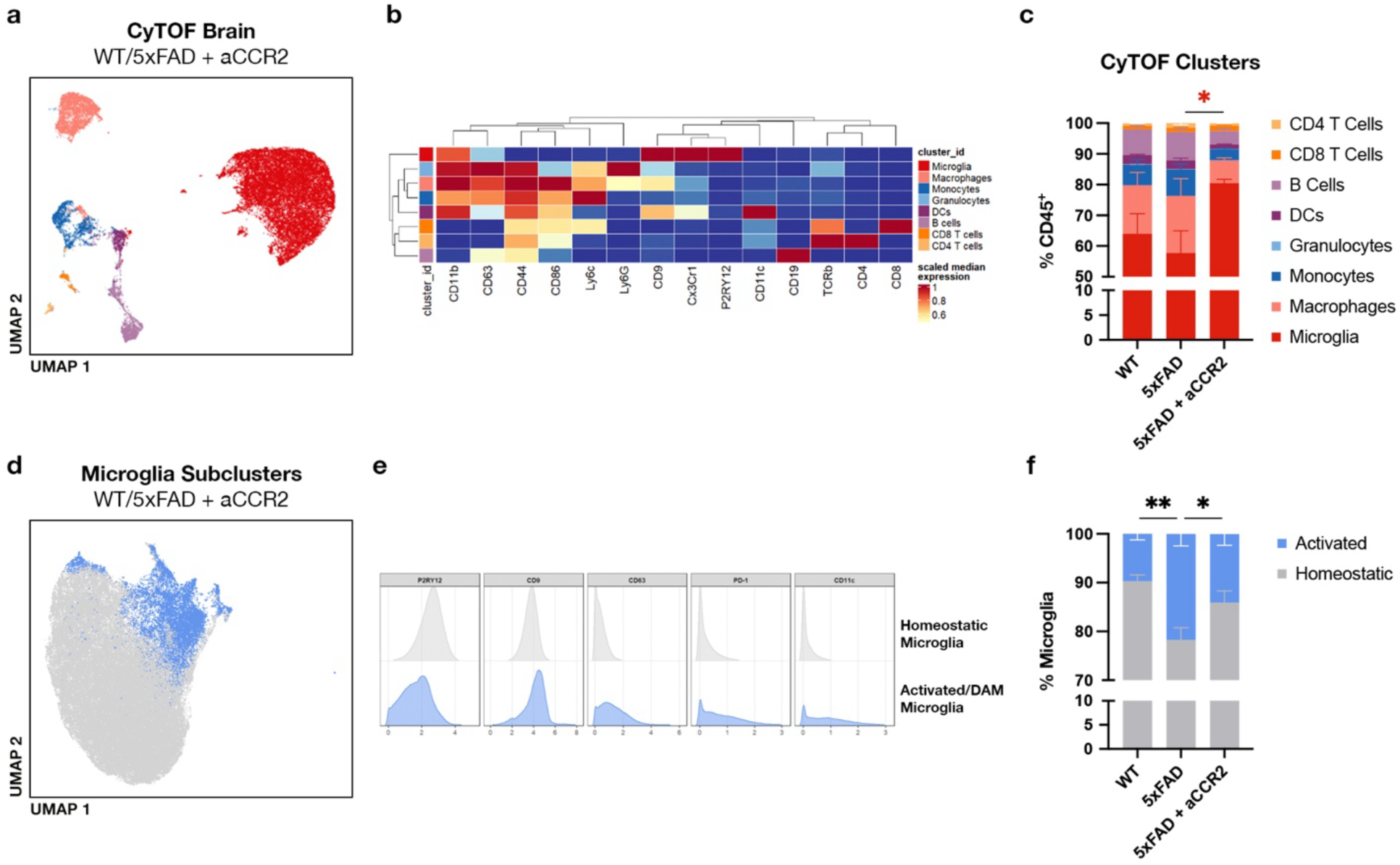
Blockade of the CCR2-CCL2 axis reduces monocyte homing to the brain and microglial activation. **a,** UMAP dimensional reduction of CD45^+^ cells from the brains of WT and 5xFAD mice treated with anti-CCR2 blocking antibody. **b**, Heatmap showing immune markers used in the CyTOF analysis to annotate the immune clusters. **c**, Immune cell proportions in the brains of WT and 5xFAD mice treated with anti-CCR2. **d**, Subclustering of microglial cells from the brains of WT and 5xFAD mice treated with anti-CCR2. Clusters of activated microglia are highlighted in blue. **e**, Histograms showing the expression of different activation and homeostatic microglial markers in the two annotated subpopulations. **f**, Proportions of activated or homeostatic microglia in the brains of WT and 5xFAD mice treated with anti-CCR2. One-way ANOVA followed by Dunnett’s multiple comparisons test. Data are represented as mean ±SEM. *p < 0.05, **p < 0.01

**Extended Data Fig. 5:**
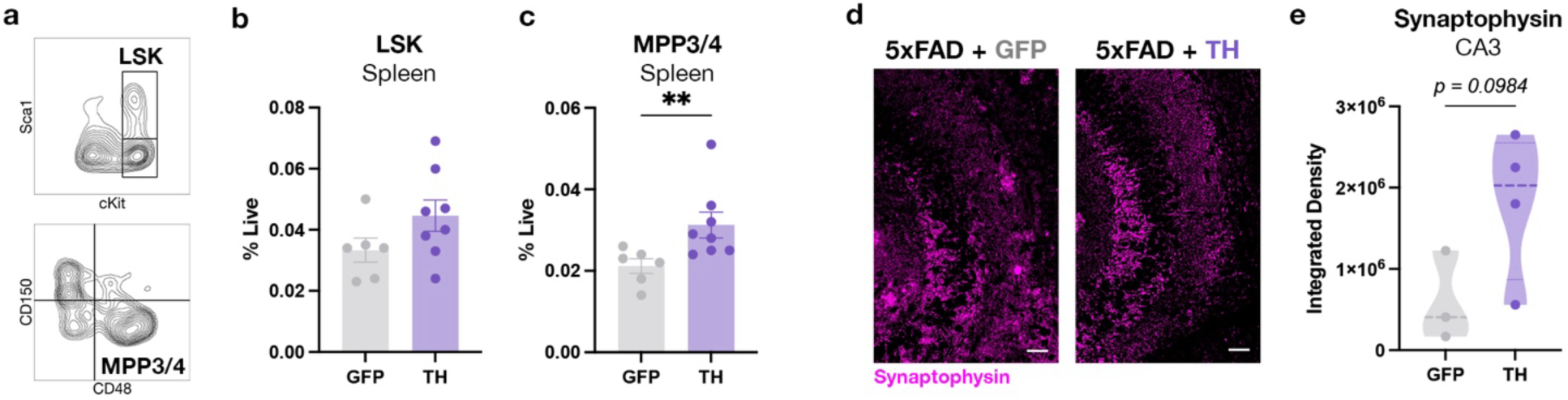
Tyrosine hydroxylase overexpression in the spleen promotes extramedullary hematopoiesis and synaptic recovery in the 5xFAD model. **a**, Flow cytometry gating strategy of HSPC subsets in the spleen of 5xFAD mice. **b**, Abundance of Lin^-^Sca1^+^cKit^+^ (LSK) hematopoietic progenitors in the spleen of 5xFAD mice overexpressing GFP or tyrosine hydroxylase (TH). **c**, Multipotent progenitors with lymphoid and myeloid potential (MPP3/4) in the spleen of 5xFAD mice overexpressing GFP or TH. **d**, Representative images of synaptophysin staining in the CA3 region of the hippocampus of 5xFAD mice overexpressing GFP or TH. Scale bar: 50 μm. e, Synaptophysin quantification in the CA3 region of the hippocampus of 5xFAD mice overexpressing GFP or TH. Unpaired t-test. In all panels, data are represented as mean ±SEM. **p < 0.01.

**Supplementary Table 1.** Flow Cytometry antibodies.

| **Antigen** | **Fluorophore** | **Clone** | **Supplier** | **Ref. No.** | **Dilution** |
| --- | --- | --- | --- | --- | --- |
| CD117 (cKit) | PE | 2B8 | BioLegend | 105808 | 1/200 |
| CD11b | PE | M1/70 | BioLegend | 101207 | 1/500 |
| CD11b | FITC | M1/70 | BioLegend | 101206 | 1/500 |
| CD150 | PE-Cy5 | TC15-12F12.2 | BioLegend | 115912 | 1/200 |
| CD44 | APC | IM7 | BioLegend | 103011 | 1/200 |
| CD45 | BV421 | 30-F11 | BioLegend | 103134 | 1/200 |
| CD48 | APC-Cy7 | HM48-1 | BioLegend | 103432 | 1/200 |
| Lineage Cocktail | Biotin | --- | Miltenyi Biotec | 130-092-613 | 1/20 |
| Ly-6C | PE-Dazzle | HK1.4 | BioLegend | 128044 | 1/200 |
| Ly-6G | APC | 1A8 | BioLegend | 127613 | 1/500 |
| Ly-6A/E (Sca-1) | PE-Cy7 | D7 | BioLegend | 108114 | 1/200 |
| Streptavidin | FITC | --- | BioLegend | 405202 | 1/200 |
| TCRbeta | APC-Cy7 | H57-597 | BioLegend | 109220 | 1/200 |

**Supplementary Table 2.**
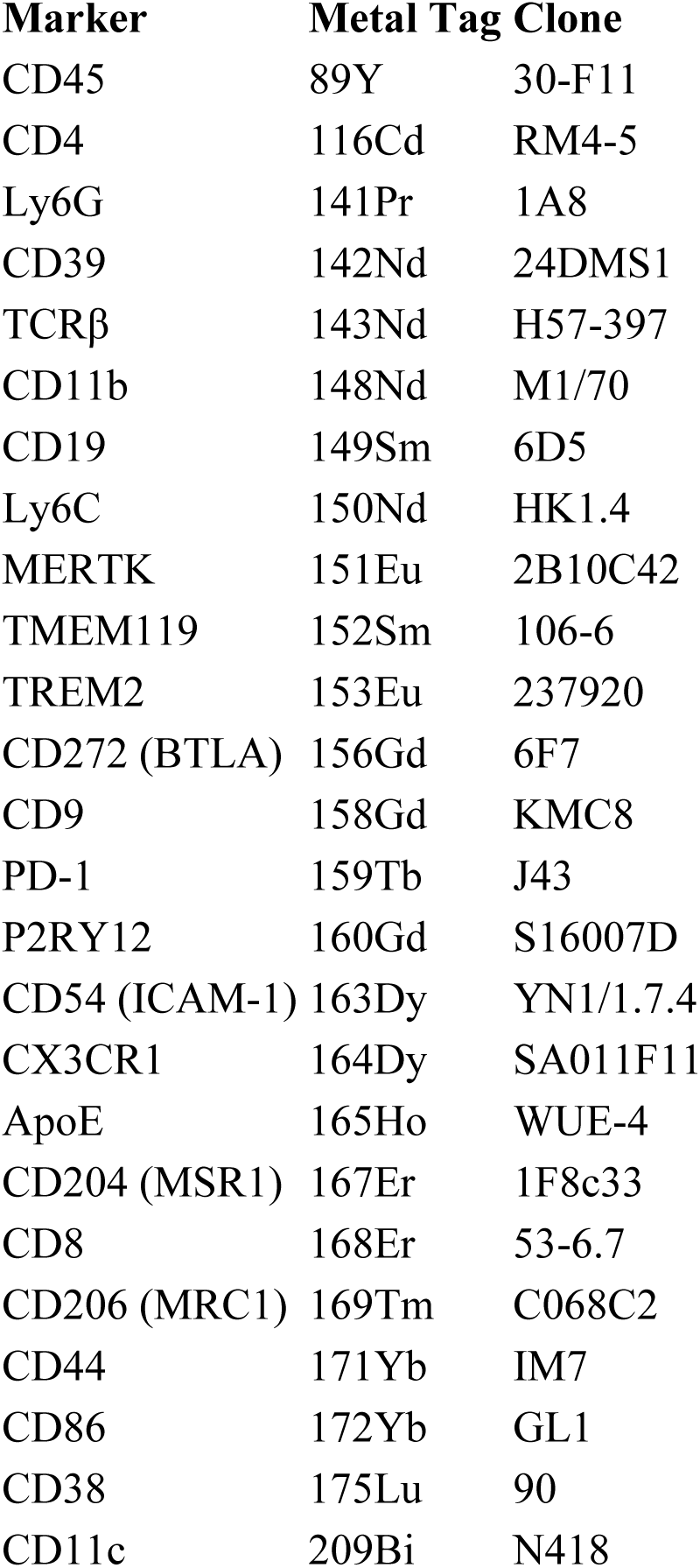
CyTOF pannel (all concentrations were used in 1:100)

